# Human axial progenitors generate trunk neural crest cells

**DOI:** 10.1101/272591

**Authors:** Thomas J. R. Frith, Ilaria Granata, Erin Stout, Matthew Wind, Oliver Thompson, Katrin Neumann, Dylan Stavish, Paul R. Heath, James O.S. Hackland, Konstantinos Anastassiadis, Mina Gouti, James Briscoe, Val Wilson, Mario R. Guarracino, Peter W. Andrews, Anestis Tsakiridis

**Affiliations:** Centre for Stem Cell Biology, Department of Biomedical Science, The University of Sheffield, Alfred Denny Building, Western Bank, Sheffield S10 2TN United Kingdom; Computational and Data Science Laboratory (CDS-LAB), High Performance Computing and Networking Institute (ICAR), National Research Council of Italy (CNR), Via Pietro Castellino, 111, Napoli, Italy; Stem Cell Engineering, Biotechnology Center, TU Dresden, Tatzberg 47, 01307 Dresden, Germany; Sheffield Institute for Translational Neuroscience, University of Sheffield, 385a Glossop Road, Sheffield S10 2HQ United Kingdom; Max Delbrück Center for Molecular Medicine, Robert-Rössle-Strasse 10, 13125 Berlin, Germany; The Francis Crick Institute, 1 Midland Road, London, NW1 1AT United Kingdom; MRC Centre for Regenerative Medicine, Institute for Stem Cell Research, School of Biological Sciences, University of Edinburgh, 5 Little France Drive, Edinburgh EH16 4UU United Kingdom

## Abstract

The neural crest (NC) is a multipotent embryonic cell population generating distinct cell types in an axial position-dependent manner. The production of NC cells from human pluripotent stem cells (hPSCs) is a valuable approach to study human NC biology. However, the origin of human trunk NC remains undefined and therefore current in vitro differentiation strategies induce only a modest yield of trunk NC cells. Here we show that hPSC-derived axial progenitors, the posteriorly-located drivers of embryonic axis elongation, give rise to trunk NC cells and their derivatives. Moreover, we define the molecular signatures associated with the emergence of human NC cells of distinct axial identities in vitro. Collectively, our findings indicate that there are two routes toward a human post-cranial NC state: the birth of cardiac and vagal NC is facilitated by retinoic acid-induced posteriorisation of an anterior precursor whereas trunk NC arises within a pool of posterior axial progenitors.

## Introduction

The neural crest (NC) is a multipotent cell population which arises in the dorsal neural plate/non-neural ectoderm border region during vertebrate embryogenesis. Studies utilising chick and amphibian embryos have indicated that different levels of BMP, WNT and FGF signals, emanating from the mesoderm/non-neural ectoderm, orchestrate NC induction and specification (1). This occurs via the action, first of neural plate border-specific transcription factors such as PAX3/7, MSX and ZIC family members, and then via definitive NC-specifiers (e.g. SOX9/10) (2). Once specified, NC cells undergo epithelial-to-mesenchymal transition (EMT), exit the neural tube, and migrate to generate various cell types. The identity of NC products correlates to their position along the anteroposterior (A-P) axis, which is in turn reflected by the expression of paralogous HOX gene groups. Cranial NC cells give rise to mesoectodermal derivatives (e.g. dermis, cartilage, bone), melanocytes, neurons and glia colonizing the head (3) and are divided into an anterior HOX-negative and a posterior HOX group(1-3)-positive domain. The latter also includes cells contributing to heart structures (termed cardiac NC) (3, 4). Vagal NC cells, which are located between somites 1-7, are marked by the expression of HOX group(3-5) members (5-7) and generate the enteric nervous system (ENS) (8). HOX group(5-9)-positive NC cells at the trunk level (5, 9-11) produce sympathoadrenal cells, which in turn give rise to sympathetic neurons, neuroendocrine cells, and melanocytes (12).

An attractive approach for studying human NC biology and modelling NC-associated developmental disorders (neurocristopathies) involves the in vitro differentiation of human pluripotent stem cells (hPSCs) toward NC cells. Conventional protocols to obtain NC from hPSCs are conceptually based on the production of a neurectodermal intermediate, via TGFβ signalling inhibition, which is subsequently steered toward a NC fate, usually through stimulation of WNT activity combined with the appropriate BMP level (13-16). These strategies yield NC cells of an anterior cranial character lacking HOX gene expression and the generation of more posterior HOX+ NC subtypes typically relies on the addition of retinoic acid (RA) and/or further WNT signalling stimulation (17-20). However, these signals fail to efficiently induce a high number of NC cells of a HOX(5-9)+ trunk identity from an anterior cranial progenitor. Therefore, the generation of trunk NC derivatives such as sympathoadrenal cells often requires the flow cytometry-based purification of small cell populations positive for lineage-specific fluorescent reporter (18) or cell surface markers (21), a time-consuming and laborious approach.

A number of studies in chicken and mouse embryos employing both fate mapping and lineage tracing have shown the existence of a posterior NC progenitor entity, which is distinct from its more anterior counterparts and potentially co-localises with a pool of caudally-located axial progenitors (22-27). These progenitors include a bipotent stem cell-like population that fuels embryonic axis elongation through the coordinated production of spinal cord neurectoderm and paraxial mesoderm (PXM) (28) (reviewed in (29) and (30). In both mouse and chick embryos these neuromesodermal progenitors (NMPs) are located in the node/streak border and the caudal lateral epiblast during early somitogenesis, and later in the chordoneural hinge within the tailbud (TB) (27, 31-34). No unique NMP markers have been determined to date and thus phenotypically NMPs are defined by the co-expression of the pro-mesodermal transcription factor Brachyury (T) and neural regulator SOX2 (35-37). Furthermore, they express transcripts which are also present in the primitive streak (PS) and TB marking committed PXM and posterior neurectodermal progenitors such as *Cdx* and *Hox* gene family members, *Tbx6* and *Nkx1-2* (23, 24, 31, 38, 39). T and SOX2 have a critical role, in conjunction with CDX and HOX proteins, in regulating the balance between NMP maintenance and differentiation by integrating inputs predominantly from the WNT and FGF signalling pathways (27, 38-41). The pivotal role of these pathways has been further demonstrated by recent studies showing that their combined stimulation results in the robust induction of T+SOX2+ NMP-like cells from mouse and human PSCs (42-44).

NMPs/axial progenitors appear to be closely related to trunk NC precursors in vivo. Specifically, trunk NC production has been shown to be controlled by transcription factors which also regulate cell fate decisions in axial progenitors such as CDX proteins (45-47) and NKX1-2 (48). The close relationship between bipotent axial and posterior NC progenitors is further supported by fate mapping experiments involving the grafting of a portion of E8.5 mouse caudal lateral epiblast T+SOX2+ cells (27) and avian embryonic TB regions (22, 34) as well as lineage tracing experiments (23, 24) which have revealed the presence of cell populations exhibiting simultaneously mesodermal, neural and NC differentiation potential. Together these findings suggest that the trunk/lumbar NC is likely to originate from a subset of axial progenitors arising near the PS/TB.

Here we sought to determine whether trunk NC is also closely related to axial progenitors in the human and thus define a robust improved protocol for the production of trunk NC cells and their products from hPSCs. We show that hPSC-derived, “pre-neural” axial progenitors contain a subpopulation that displays a BMP-dependent, mixed early NC/NMP transcriptional signature and thus is likely to represent the earliest trunk NC precursors. We demonstrate that neuromesodermal-potent axial progenitor cultures are competent to generate efficiently trunk NC cells, marked by thoracic HOX gene expression. This transition to trunk NC appears to take place via the maintenance of a CDX2/posterior HOX-positive state and the progressive amplification of an NC gene regulatory network. We also show that “caudalisation” via RA treatment of anterior NC precursors leads to the acquisition of a mixed cardiac and vagal NC identity rather than a trunk NC character and define novel markers of distinct posterior NC subtypes. Finally, we utilise our findings to establish a protocol for the in vitro generation of PHOX2B+ sympathoadrenal cells and sympathetic neurons at high efficiency from cultures of posterior axial progenitor-derived trunk NC cells without the need for FACS-sorting to select for minor precursor subpopulations. Taken together these findings provide an insight into the mechanisms underpinning the “birth” of human NC cells at different axial levels and pave the way for the modelling of trunk NC-linked diseases such as neuroblastoma.

## Results

### Transcriptome analysis of human axial progenitors

We and others have previously shown that combined stimulation of the WNT and FGF signalling pathways in PSCs leads to the production of a high (>80%) percentage of T+SOX2+ cells. The resulting cultures resemble embryonic posterior axial progenitors, including NMPs, both in terms of marker expression and developmental potential (38, 42-44, 49). To interrogate the transcriptome changes associated with the induction of such progenitors in a human system and identify the presence of trunk NC precursors, we carried out RNA sequencing (RNAseq) following 3-day treatment of hPSCs with recombinant FGF2 and the WNT agonist/GSK-3 inhibitor CHIR99021 (CHIR). As reported previously, most cells emerging under these conditions co-expressed T and SOX2 as well as CDX2 (**Figure 1A**). We found that the transcriptomes of hNMPs and hPSCs were markedly distinct from each other (**Figures S1A, S1B**) and dramatic global gene expression changes accompanied the acquisition of an axial progenitor character: 1911 and 1895 genes were significantly (padj<0.05; Fold Change >=2) up- and down-regulated compared to hPSCs respectively (**Table S1**). Predictably, the top-downregulated genes were associated with undifferentiated hPSCs (e.g. NANOG, GDF3, POU5F1), anterior neurectoderm (OTX2) and lateral/ventral mesoderm (KDR). The vast majority of the top-upregulated genes were well-established drivers of axis elongation (e.g. *TBRA*, *CDX1/2*, *EVX1*, *MSGN1*, *TBX6*) and WNT/FGF/NOTCH/RA signalling pathway components, known to be expressed at high levels in the late PS/TB regions in vivo (e.g. *WNT3A/5B*, *RSPO3*, *FGF4/8*, *FGF17*, *HES7*) (**Figures 1B**, **S1C, S1D, Table S1**). A large fraction of upregulated genes were transcriptional regulators (**Figure S1D**, **Table S1**) and we found that members of HOX gene clusters belonging to paralogous groups 1-9 were strongly differentially expressed between the two groups (**Figure 1C**, **S1E, Table S1**). The upregulation of posterior thoracic (group 5-9) HOX transcripts as well as the presence of many transcripts (23/32) marking “late” E9.5 mouse embryonic NMPs such as CYP26A1, FGF17 and WNT5A (38) suggest that day 3 WNT-FGF-treated hPSC cultures may correspond to a more developmentally advanced axial progenitor state. Overall, these data confirm our previous observations that treatment of hPSCs with WNT and FGF agonists gives rise to cultures resembling embryonic posterior axial progenitors.

**Figure 1.**
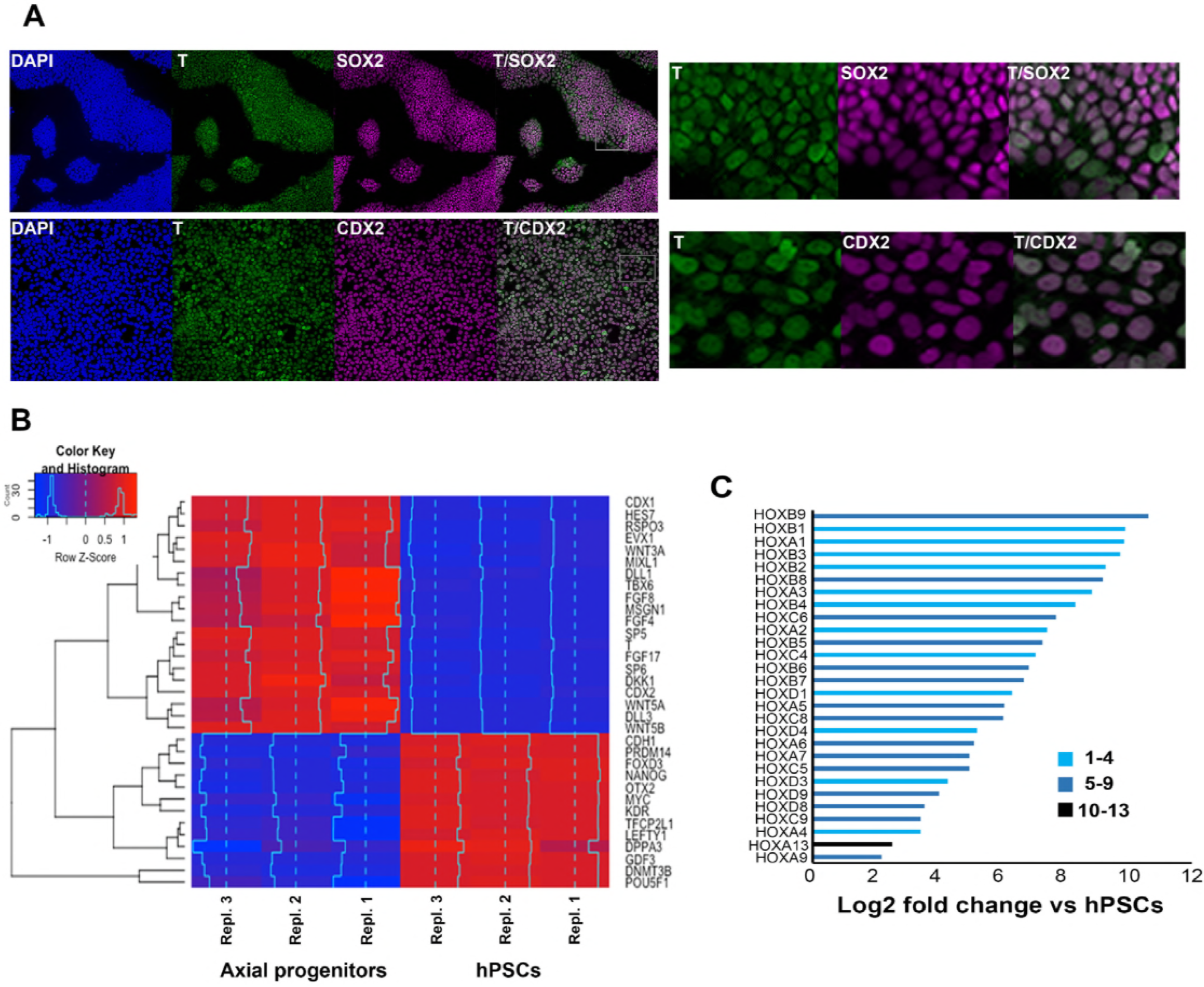
Transcriptome analysis of in vitro-derived human axial progenitors. (A) Immunofluorescence analysis of expression of indicated markers in day 3 hPSC-derived axial progenitors. Magnified regions corresponding to the insets are also shown. (B) Heatmap showing the expression values of selected markers in three independent axial progenitor and hPSC sample replicates. The expression values (FPKM) were scaled to the row mean. The color key relates the heat map colors to the standard score (z-score). (C) Induction of all significantly upregulated HOX transcripts in axial progenitors relative to hPSCs. Paralogous HOX groups corresponding to different axial levels such as cervical (groups 1-4), brachial/thoracic (5-9) and lumbosacral (10-13) are indicated. See also Figure S1 and Table S1.

### A BMP-dependent neural crest signature in human axial progenitor cultures

We next sought evidence pointing to the potential links between trunk NC and human axial progenitors. The RNAseq analysis revealed that a considerable number of genes known to collectively mark the neural plate border and early NC in vivo (“NC/border” e.g. SOX9, PAX3, MSX1/2, SNAI1/2, ZIC1/3) were also significantly upregulated in axial progenitors (**Figure 2A**), a finding which was further verified using quantitative real time PCR (qPCR) (**Figure S2A**). To exclude the possibility that the presence of such markers was the result of spontaneous differentiation of NM bipotent axial progenitors and their neural derivatives we examined their co-expression with T, a marker of both NMPs and prospective PXM. Immunostaining of d3 WNT-FGF-treated hPSC cultures showed that a considerable fraction of T+ cells also expressed the early NC markers/specifiers SOX9, SNAI2 and PAX3 (**Figure 2B**). Moreover, SOX9-positive cells were found to express low or no MSGN1 or TBX6 (**Figure S2B**) suggesting that the upregulation of border/NC markers following WNT-FGF treatment of hPSCs was unlikely to reflect the presence of committed PXM cells given that some of these genes are also expressed in the mesoderm during axis elongation. Collectively, these findings indicate that a NC/border state probably arises within multipotent posterior axial progenitors which have not committed to a neural or mesodermal fate.

A number of both in vivo and in vitro studies have pointed to an optimal level of low/intermediate BMP signalling acting as an inducer of a NC/border character in conjunction with WNT and FGF (14, 16, 48, 50-53). We therefore examined whether hPSC-derived axial progenitor cultures exhibit endogenous BMP activity. RNAseq revealed that many BMP pathway-associated transcripts (BMP2/4/6/7 and ID1/4) were significantly upregulated compared to hPSCs (**Figure 2A**). Moreover, antibody staining showed expression of phosphorylated SMAD1/5, a readout of BMP activity (**Figure 2C**). This is extinguished upon treatment with the BMP antagonist LDN193189 (LDN) (54) during the in vitro differentiation of hPSCs to day 3 axial progenitors (**Figure 2C**). Interestingly, BMP inhibition also caused a decrease in the transcript levels of most border/NC-specific transcripts with PAX3 and MSX1 being the most severely affected (**Figure 2D**). We also observed a reduction in the NMP/late PS/TB markers CDX2 and WNT3A whereas the PXM specifier TBX6 remained relatively unaffected by LDN treatment suggesting that emergence of prospective PXM is not influenced by BMP inhibition (**Figure 2D**). By contrast, SOX2 expression was markedly increased upon LDN treatment (**Figure 2D**). Taken together these data indicate a correlation between endogenous BMP activity and the acquisition of a SOX2+low border/NC identity by posterior axial progenitors while transition toward a SOX2+ high neural fate relies on BMP antagonism in vitro.

**Figure 2.**
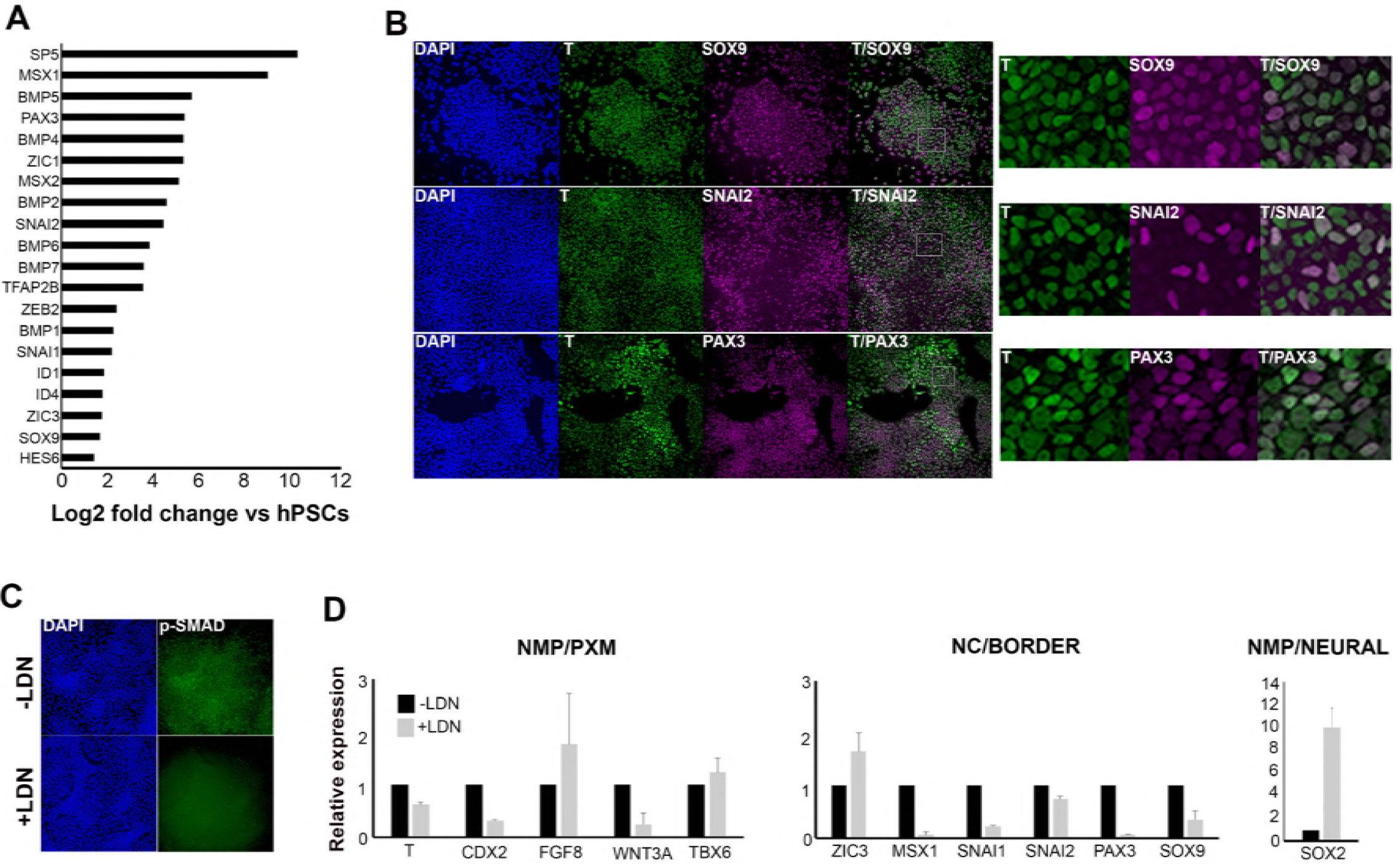
hNMP cultures exhibit a BMP-dependent neural crest/border signature. (A) Log-fold induction of representative neural crest/neural plate border and BMP-associated transcripts in axial progenitors compared to hESCs. (B) Immunofluorescence analysis of expression of indicated markers in axial progenitos. Magnified regions corresponding to the insets are also shown. (C) Immunofluorescence analysis of expression of phosphorylated SMAD1/5 (p-SMAD) in the presence and absence of the BMP inhibitor LDN193189 (LDN). (D) qPCR expression analysis of indicated markers in axial progenitors in the presence (+) or absence (−) of LDN. Error bars=S.D. (n=2). In all cases nuclei were counterstained with DAPI. PXM, paraxial mesoderm; NC, neural crest. See also Figure S2.

### In vitro-derived axial progenitors are a source of trunk neural crest cells

We reasoned that if posterior axial progenitors with border/NC features correspond to pioneer trunk NC precursors then they should be competent to generate definitive trunk neural crest when placed in the right culture environment. We have recently reported a protocol for the efficient generation of anterior cranial NC cells from hPSCs involving the combined stimulation of WNT signalling, TGFβ signalling inhibition and moderate BMP activity via the parallel addition of BMP4 and the BMP type 1 receptor inhibitor DMH1 (16). Culture of day 3 WNT-FGF-treated hPSCs under these NC-inducing conditions for 5-6 days gave rise to a high number (average percentage=50% of total cells) of cells co-expressing the definitive NC marker SOX10 together with HOXC9, a readout of trunk axial identity (**Figure 3A, 3B, 3C**). A large proportion of the cultures were also SOX9+HOXC9+ further confirming an trunk NC character whereas the percentage of neural cells marked by SOX1 expression remained very low throughout the course of the differentiation (**Figure 3B**, **S3A, S3B**). This may indicate that posterior NC progenitors do not progress through neural commitment but rather diverge from an earlier pre-neural, border-like stage reflecting previous reports which show that NC specification takes place prior to definitive neurulation (48, 55, 56). Furthermore, during the transition toward trunk neural crest, the NMP/pre-neural marker NKX1-2 was rapidly extinguished followed shortly after by T, while CDX1 transcript levels declined more slowly (**Figure S3A**, **S3B**). By contrast, the expression of CDX2 and SOX9 was maintained at high levels (>70% of total cells) throughout the course of differentiation of axial progenitors to trunk NC while SOX10 expression appeared only after day 7 of differentiation (Day 0 defined as the start of axial progenitor induction from hPSCs) (**Figure S3C**, **data not shown**). We also confirmed the NM potency of the starting axial progenitor cultures as treatment with high levels of FGF2/CHIR and RA led to the production of TBX6+ PXM and SOX1+ spinal cord, posterior neurectoderm (PNE) cells respectively (**Figure 3A, 3D**). Taken together these data suggest that hPSC-derived NM-potent axial progenitor cultures are competent to produce trunk NC at high efficiency.

**Figure 3.**
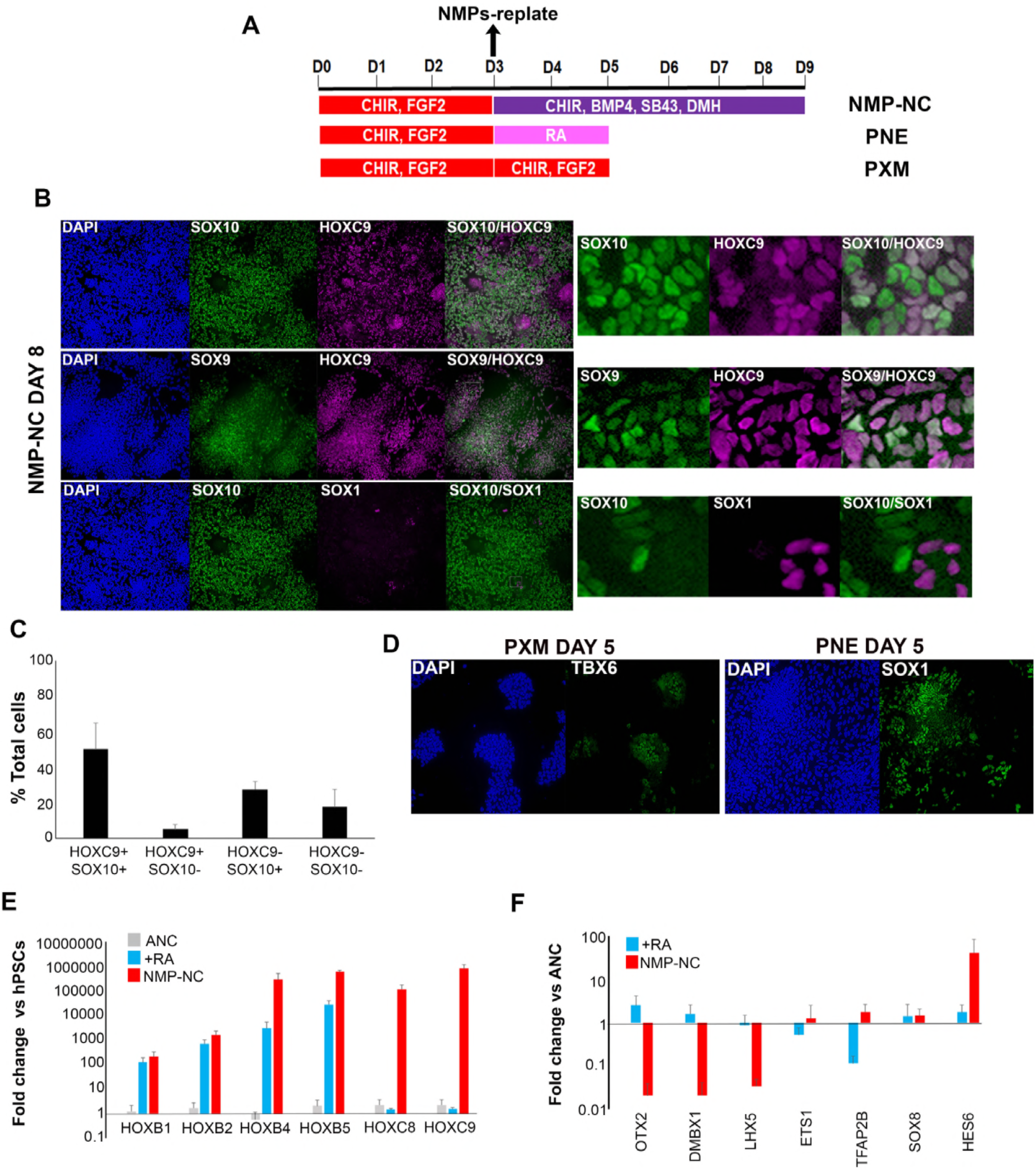
In vitro derived axial progenitors generate trunk neural crest efficiently. (A) Diagram depicting the culture conditions employed to direct trunk NC, posterior neurectoderm (PNE) and paraxial mesoderm (PXM) differentiation from hPSC-derived axial progenitors. (B) Immunofluorescence analysis of the expression of indicated NC (SOX10, SOX9) and neural (SOX1) and the thoracic/trunk marker HOXC9 in trunk NC (TNC) cells derived from axial progenitors after 8 days of differentiation. Magnified regions corresponding to the insets are also shown. Note that the HOXC9/SOX10 and SOX10/SOX1 image sets correspond to the same cells. (C) Quantitation of cells marked by different combinations of HOXC9 and SOX10 expression in day 8 trunk NC cultures derived from axial progenitors following image analysis. Six randomly selected fields (two independent experiments) were used for analysis (total number of cells scored=5366, average number of cells/field=894, error bars=s.d.). (D) Immunofluorescence analysis of TBX6 (left) or SOX1 (right) expression in axial progenitors treated with CHIR-FGF2 (pro-PXM conditions) and RA (pro-PNE conditions) respectively. (E) qPCR expression analysis of indicated HOX genes in hPSC-derived anterior cranial (ANC), retinoic acid (RA)-treated NC (+RA), and axial progenitor-derived NC cells (NMP-NC) relative to hPSCs. Error bars=S.E.M. (n=3). (F) qPCR expression analysis of indicated NC markers in +RA and axial progenitor-derived NC cells relative to untreated anterior cranial NC cells. Error bars=S.E.M. (n=3). See also Figure S3.

Similar to popular in vitro neural induction strategies, most current NC differentiation protocols aiming to generate posterior (e.g. trunk) cell populations from hPSCs rely on the caudalisation of an anterior ectodermal precursor via treatment with RA and/or WNT agonists (15, 17-20). Therefore, we compared our axial progenitor–based approach for generating trunk NC to a conventional strategy involving the generation of anterior cranial NC (ANC) precursor cells (16) followed by RA addition in the presence of WNT and BMP signalling (**Figure 4A**). The axial identity of the resulting cells was assessed by qPCR assay
of *HOX* transcripts corresponding to different levels along the A-P axis. In line with previous findings (17, 19) RA-treated cells expressed high levels of *HOX* group 1-5 members compared to untreated NC suggesting a posterior cranial and vagal/cardiac NC character (**Figure 3E**). However, efficient induction trunk of *HOXC8* and *9* transcripts was only achieved when posterior axial progenitors were employed as the starting population for NC generation (**Figure 3E**). Furthermore, axial progenitor-derived NC cells were marked by increased expression of the trunk NC marker *HES6*, but did not express the cranial markers *OTX2*, *DMBX1* and *LHX5* although they were positive for the “late” cranial NC transcripts (*TFA2B*, *ETS1*, *SOX8*) (57) (**Figure 3F**). We thus conclude that posterior axial progenitors are the ideal starting population for efficiently generating trunk NC in vitro whereas RA treatment of anterior NC precursors predominantly produces posterior cranial and cardiac/vagal NC cells. These data also serve as evidence supporting the notion that trunk NC precursors are likely to arise within cells with axial progenitor/NMP features rather than a caudalised anterior progenitor.

**Figure 4.**
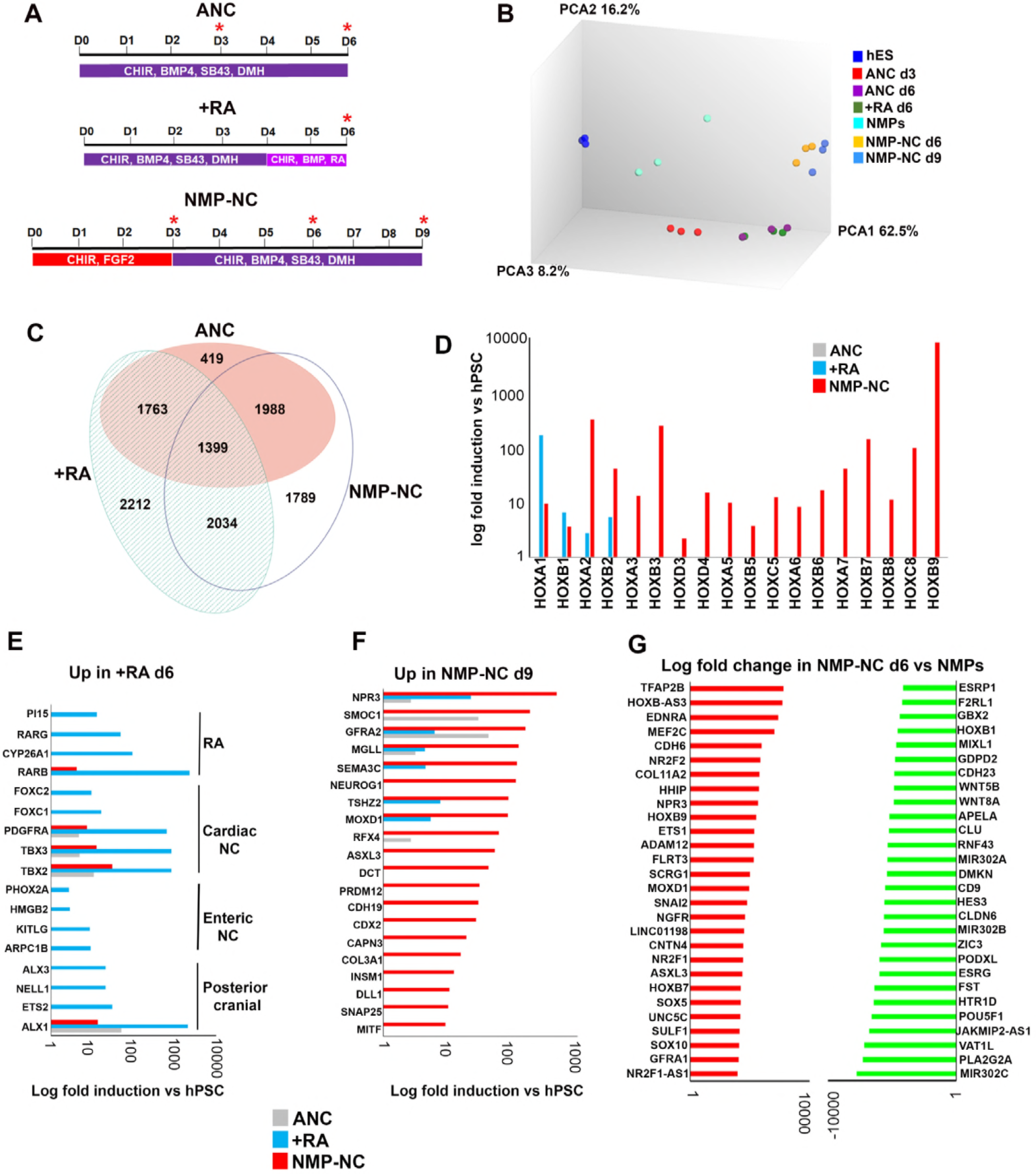
Transcriptome analysis of in vitro derived neural crest cells corresponding to distinct axial levels. (A) Diagrams showing the culture conditions employed for generating NC cells of distinct axial identities using hPSCs. Asterisks indicate the timepoints used for sample harvesting and transcriptome analysis. D, day of differentiation. ANC, Anterior neural crest. (B) Principal component analysis depicting variance between different samples used for microarray analysis (timepoints shown in A). (C) Venn diagram showing the overlap between all significantly upregulated (≥2-fold relative to undifferentiated hESCs, FDR ≤ 0.05) in each indicated NC group. (D) Log fold induction of HOX genes in indicated NC populations relative to hPSCs. (E) Log fold induction of representative significantly upregulated (≥2-fold relative to undifferentiated hPSCs, FDR ≤ 0.05) transcripts marking day 6 RA-treated NC cells. (F) Log fold induction of representative significantly upregulated (≥2-fold relative to undifferentiated hPSCs, FDR ≤ 0.05) transcripts marking day 9 axial progenitor-derived NC cells. (G) Log fold changes in the expression of the most-upregulated and most-downregulated transcripts in day 6 axial progenitor-derived NC precursors compared to d3 hPSC-derived axial progenitors. See also Figure S4, Tables S2, S3, and S4.

### Efficient A-P patterning of human neural crest cells reveals molecular signatures of distinct axial identities

To further discern the identity of posterior NC subtypes induced either via RA treatment or an axial progenitor intermediate as well as identify unique associated molecular signatures we carried out analysis of the transcriptomes of NC cells arising under these conditions as well as those of their precursors using microarrays (**Figure 4A**). We found that axial progenitor-derived NC cells (NMP-NC d9) and their precursors (NMP-NC d6) grouped together and were distinct from a cluster containing d6 anterior cranial NC (ANC) and +RA NC cells and their common d3 progenitor (ANC d3) (**Figure 4B**, **S4A**). Although the three final populations exhibited distinct transcriptional profiles they all expressed pan-NC genes including “early” NC/border (MSX1/2, PAX3/7) and “late” NC (SOX10, SNAI1/2) transcripts (**Figure 4C**, **S4B, S4C, Table S2**). In line with our previous observations (**Figure 3C**), ANC cells failed to express any HOX transcripts, RA treatment induced anterior HOX genes and only WNT-FGF-treated hPSCs gave rise to NC cells positive for thoracic HOX group (5-9) members (**Figure 4D**) reflecting an anterior cranial, posterior cranial/cardiac/vagal and trunk NC fate respectively. The axial identity of the resulting posterior NC subtypes was further confirmed by the observation that some of the most-upregulated transcripts in +RA cells were established posterior cranial (e.g. ALX1/3 (58)), cardiac (e.g. *FOXC1* and 2 (59); *PDGFRa* (60); *TBX2/3* (61)) and vagal/enteric NC markers (*PHOX2A* (62); *KITLG* (63); *ARPC1B* (64) (**Figure 4E)**. In contrast, known trunk NC, sympathoadrenal and sympathetic/sensory neuron regulators such as CDX2 (47), *INSM1* (65), *ISL1* (66) and *NEUROG1* (67) were induced only in NC cells derived from axial progenitors (**Figure 4F**, **Table S2**). We also identified *ASLX3*, the human homologue of the Drosophila polycomb protein *asx* (68) which has been recently linked to the developmental syndrome Bainbridge-Ropers (69) as a novel trunk NC marker (**Figure 4F**, **Table S2**). Transcription factors specifically induced in anterior cranial NC cells included the forkhead gene *FOXS1* which has been shown to be expressed in mouse NC derivatives (70) and *TCF7L2*, a WNT signalling effector which has been reported to harbour a NC-associated enhancer (71) (**Tables S2, S3**). Collectively these data support the idea that a mixed posterior cranial/vagal/cardiac NC character arises upon treatment of anterior NC precursors with RA whereas a *bona fide* trunk NC identity can be achieved only via an axial progenitor intermediate.

One of the most over-represented gene categories in all three axial NC subtypes were transcription factors and a common NC-specific transcription factor module was found to be expressed regardless of axial character (**Figure S4B**, **Table S3, Table S4**). This included well-established border/NC regulators such as *PAX3/7*, *MSX2*, *SOX9/10*, *TFAP2A-C* and *SNAI1/2* (**Figure S4C**, **Tables S3, S4**). However, the expression levels of many of these transcription factors varied between the three groups (**Figure S4C**). The highest levels of HES6, FOXD3 and MSX1/2 were found in axial progenitor-derived trunk NC cells whereas high PAX7 and SNAI1/ SOX9 expression was more prevalent in the anterior cranial and RA-treated samples respectively (**Figure S4C**). Comparison of the day 6 trunk and d3 ANC precursor transcriptomes also revealed that expression of LHX5 and DMBX1 marks an anterior NC state whereas HES6 is associated exclusively with a trunk fate (**Figure S4D**) indicating that diversification of axial identity in NC cells starts at an early time point via the action of distinct molecular players.

### Distinct routes to posterior neural crest fates

To identify candidate genes mediating the gradual lineage restriction of trunk NC precursors present in axial progenitor cultures we compared the transcriptomes of d6 trunk NC precursors and day 3 WNT-FGF-treated hPSCs (=”NMPs”). We found that dramatic global gene expression changes take place during the axial progenitor-trunk NC transition (**Figure 4G**, **S4E**). Some of the most upregulated transcripts were the NC-specific TFAP2A/B, ETS1, SOX5 and SOX10 together with the established trunk NC specifier CDX2, the novel trunk NC marker ASLX3, the nuclear receptors NR2F1/2 and thoracic HOX genes (HOXB7, B9) (**Figure 4G**, **Table S4**). In contrast, signature axial progenitor transcription factors such as *T*, *MIXL1*, *NKX1-2*, anterior HOX genes (*HOXA1/B1*) and some WNT signalling components (*WNT8A/5B*) were significantly downregulated (**Figure 4G**, **Table S4**). Thus differentiation of trunk NC precursors appears to involve the transition from an axial progenitor-associated gene regulatory network to a NC-specifying one that incorporates factors which potentially act as general determinants of posterior cell fate (*CDX2, HOXB9*).

We also examined transcriptome changes during the transition from an anterior NC precursor state (ANC d3) to RA-posteriorised vagal/cardiac NC cells (+RA d6). The most-highly induced transcripts in posterior cranial/cardiac/vagal NC cells included the RA receptors beta and gamma (*RARb/g*) which have been involved in hindbrain and neural crest patterning (72) and the T-box transcription factor *TBX2*, a marker of cardiac NC and in vitro derived vagal/enteric NC progenitors (19) (**Table S4**). Other upregulated transcripts included the planar cell polarity (PCP) component *PRICKLE1*, a regulator of cardiac NC cell function (73) and the TGFβ signalling-associated gene *TGFBI* (**Table S4**). Anterior NC d3 precursor specific–transcripts included the border markers *PAX7* and *ZIC3* as well as the early cranial NC transcription factors *OTX2* and *LHX5* (57) (**Table S4**). These results indicate that, in contrast to trunk NC cells, posterior crania/cardiac/vagal NC cells arise from an anterior neural plate border precursor through posteriorisation under the influence of RA and possibly the non-canonical WNT and TGFβ pathways.

### Efficient in vitro generation of sympathoadrenal cells from axial progenitors

We next sought to determine whether trunk NC cells derived from axial progenitors are the optimal source of sympathoadrenal (SA) progenitors and their derivatives. The BMP and sonic hedgehog (SHH) signalling pathways have been shown to be critical for the specification of these lineages from NC cells (74, 75). Therefore we cultured d8 trunk NC cells generated from axial progenitors carrying a GFP reporter within the SA/sympathetic neuron regulator PHOX2B locus (18, 76), in the presence of BMP and sonic hedgehog (SHH) signalling agonists (**Figure 5A**). GFP expression was assayed after 4 days of culture of trunk NC cells in BMP4 and SHH agonists (i.e. day 12 of differentiation) (**Figure 5B**). FACS analysis revealed that the majority of cells were PHOX2B expressing (average percentage from four independent experiments=73.5%, s.d.=6.344) (**Figure 5B**) and a large proportion of them were also positive for the early SA progenitor marker ASCL1 (77) indicating that they had acquired a symphathoadrenal identity (**Figure S5A**). Further maturation of the resulting SA progenitors in the presence of neurotrophic factors (BDNF, GDNF and NGF) for a further 6-8 days resulted in the induction of a high yield of sympathetic neurons/progenitors co-expressing PHOX2B together with the sympathetic neuron regulator GATA3 (78) (average of 40% of total cells), ASCL1 (63%) and the SA differentiation regulator *ISL1* (64%) (**Figure 5C, 5D**). A high proportion of the resulting cells also expressed the dopamine production-associated enzyme/sympathetic neuron marker tyrosine hydroxylase (TH) (79) (**Figure 5D**). Furthermore, the cultures widely expressed the peripheral nervous system marker PERIPHERIN (PRPH) (80) together with the trunk axial marker HOXC9 (**Figure S5B**). We also detected dramatic induction of *GATA3*, *ASCL1*, *TH* and *PHOX2B* transcripts (between 1000 and 1,000,000-fold) as well as other SA lineage markers such as *GATA2*, *DBH* (dopamine-β-hydroxylase) and to a lesser extent *PHOX2A* using qPCR (**Figure 5E**). Together, these results suggest that the optimal route toward the efficient production of sympathoadrenal cells and sympathetic neurons from hPSCs relies on the induction of posterior axial progenitors.

**Figure 5.**
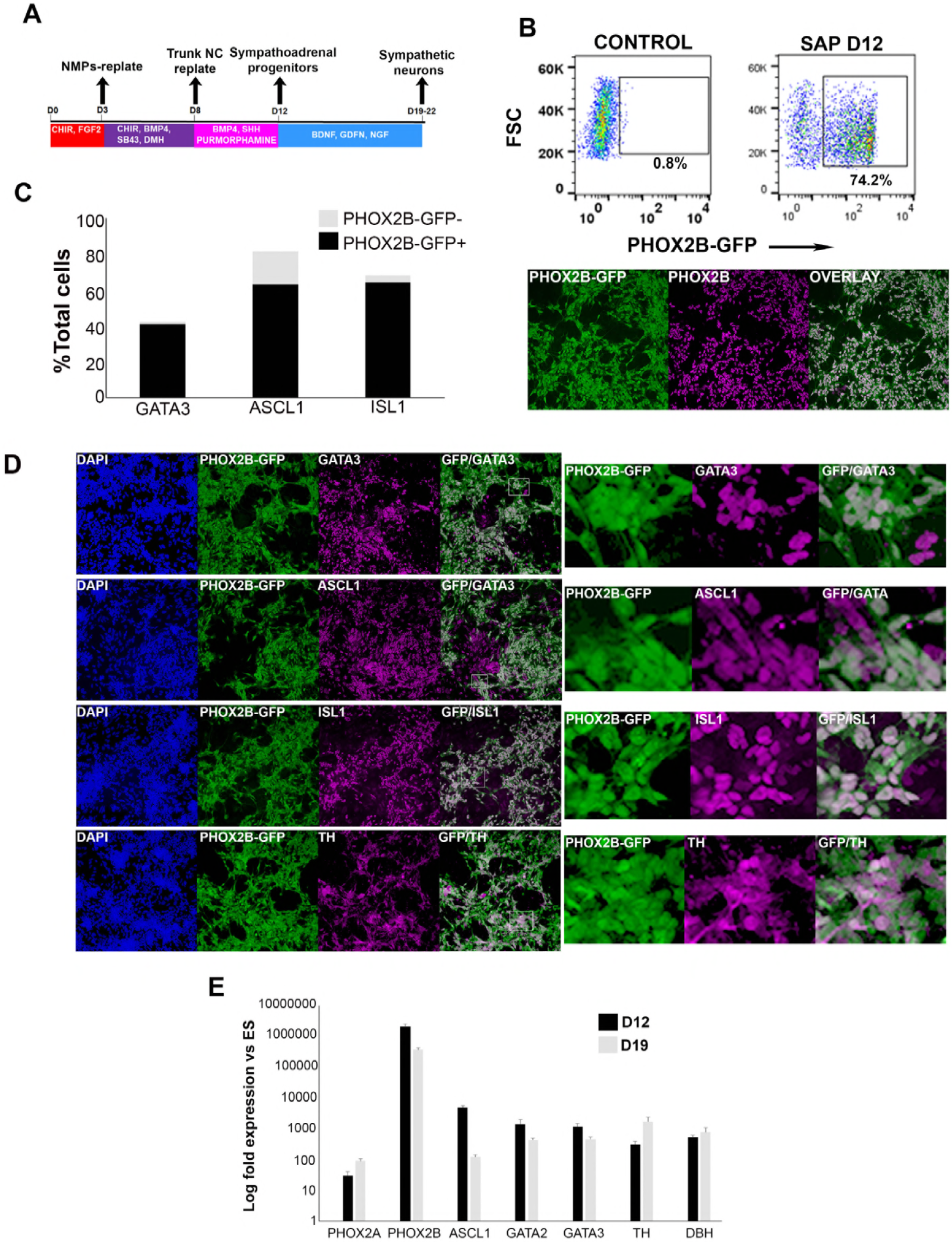
Axial progenitor-derived trunk neural crest is an optimal source of sympathoadrenal cells. (A) Diagram depicting the culture conditions employed to direct axial progenitors (“NMPs”) toward trunk NC and subsequently sympathoadrenal progenitors (SAP) and sympathetic neurons. (B) FACS analysis of PHOX2B-GFP expression in SAP cells derived from axial progenitors as shown in A. Below: Immunofluorescence analysis of PHOX2B-GFP and PHOX2B protein expression following antibody staining. (C) Quantitation of d18 differentiated cells positive for the indicated markers in relation to PHOX2B-GFP expression following antibody staining. In each case four randomly selected representative fields were used to obtain the average number of cells/marker. Total numbers of cells scored: GATA3 (N=3003), ASCL1 (N=2575), ISL1 (N=2963). (D) Immunofluorescence analysis of PHOX2B-GFP together with the indicated markers in day 18 differentiated SAP/sympathetic neurons derived from axial progenitors as shown in A. (E) qPCR expression analysis of indicated SAP/sympathetic neuron markers in d12 and d18 cultures. Error bars=S.E.M. (n=3). See also Figure S5.

## Discussion

Despite progress in the optimisation of current NC differentiation protocols the in vitro generation of trunk NC cells from hPSCs remains challenging and requires FACS-sorting of selected progenitor subpopulations, a time-consuming and laborious process also associated with increased cell death. This bottleneck prevents the dissection of the mechanisms directing human NC emergence at different axial levels as well as the efficient isolation of cell types for modelling trunk NC-specific neurocristopathies such as neuroblastoma. Previous work in amniote embryos suggested that posterior (trunk/lumbosacral) NC cells arise independently from their anterior counterparts, within a pool of axial progenitors localised near the primitive streak and the tailbud during axis elongation (22, 25-28). Here we utilised these findings and exploited our ability to induce T+ NM-potent axial progenitors from hPSCs in order to use them as the optimal starting point for the efficient in vitro derivation of trunk NC (~50% HOXC9+SOX10+), SA progenitors (~70% PHOX2B-GFP+) and sympathetic neurons without the use of FACS sorting. This strategy represents a considerable improvement over current approaches, which typically yield 5-10% PHOX2B-GFP+ cells (18) and is in line with a recent study reporting the successful production of chromaffin-like cells through the use of an NC-induction protocol which transiently produces T+SOX2+ cells (20, 21).

We show that, similar to neural cells a HOX-positive posterior identity is acquired by human NC cells via two distinct routes: posterior cranial/vagal/cardiac HOX(1-5)+ NC cells emerge through the RA/WNT-induced posteriorisation of a default anterior precursor, reflecting Nieuwkoop’s “activation-transformation” model, whereas HOX(5-9)+ trunk NC cells arise from a separate WNT/FGF-induced posterior axial progenitor exhibiting caudal lateral epiblast/NMP features mixed with a BMP-dependent neural plate border/neural crest identity (**Fig. 6**). This finding offers an explanation for the failure of current RA posteriorisation-based in vitro differentiation protocols (17, 19) to yield high numbers of HOX9+ trunk NC cells and should serve as the conceptual basis for the design of experiments aiming to generate NC cells of a defined A-P character from hPSCs.

**Figure 6.**
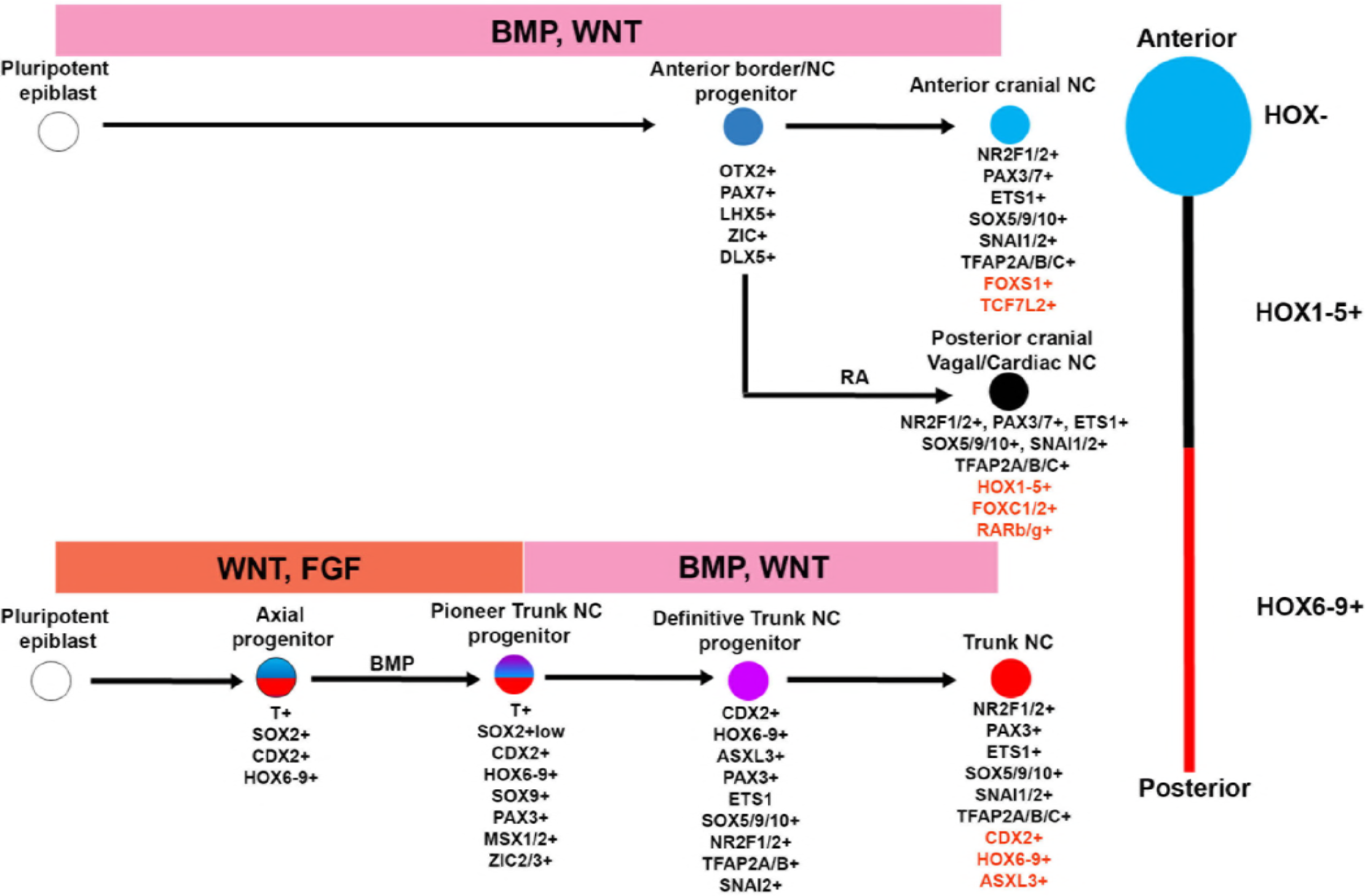
A-P patterning of in vitro derived human NC cells. Diagrammatic model summarising our findings on the in vitro generation of NC subtypes of distinct A-P identity from hPSCs. Examples of unique genes that were found to mark each NC population exclusively are shown in red.

Our data indicate that a subpopulation of in vitro derived human axial progenitors acquires border/early NC characteristics in response to the WNT and FGF signals present in the differentiation culture media, and under the influence of autocrine BMP signalling. This is in line with bulk and single cell transcriptome data showing that mouse embryonic axial progenitors/NMPs express border and early NC markers (38, 41). Furthermore, our data reflect findings in the chick embryo showing that an “unstable”, pre-neural plate border domain, potentially defined by the co-localisation of pre-neural (*Nkx1-2)* (81) and border markers such as *Pax3* (82) and *Msx1,* arises in the avian embryonic caudal lateral epibllast in response to autocrine BMP, FGF (51) and possibly WNT signalling (83). We also found that CDX2 expression is maintained at high levels during the generation of trunk NC indicating that this transcription factor might be critical in inducing an NC character in axial progenitors. CDX2 has been shown, together with β-catenin, to bind and activate neural plate border/early NC specifiers such as MSX1 and ZIC1 (47, 84) and, intriguingly, ChiP-Seq data from in vitro-derived mouse NMPs have revealed that many border/NC genes are direct targets of CDX2 often jointly with T (e.g. PAX3, SOX9, ZIC3) (39). Collectively these findings raise the possibility that β-catenin and CDX2, in conjunction with FGF/BMP signalling, may be critical for the establishment of a border/NC identity in T+ axial progenitors and further work is required to test this hypothesis.

We show that BMP/WNT treatment of human axial progenitors promotes the induction of a definitive trunk NC state. This transition appears to coincide with the progressive extinction of key axial progenitor genes and their replacement by a battery of NC-specific transcription factors such as TFAP2B, SOX10, NR2F2 and NR2F1 while the levels of some “common” axial progenitor-NC markers (e.g. SOX9, PAX3, TFAP2A and SNAI2) remain high (**Figure 4**, **S3, Table S4**). TFAP2A has been previously reported to act as a master NC transcription factor whose binding on key enhancers, together with NR2F1/2, appears to initiate transcription of NC-specific genes (71). This finding raises the possibility that these transcription factors are the molecular drivers of the transition from an early posterior axial progenitor state to a lineage-restricted trunk NC fate in response to BMP/WNT. How is a trunk axial character specified? Our transcriptome analysis data suggest that a generic trunk identity is first installed during the emergence of multipotent CDX2+trunk HOX+ axial progenitors under the influence of WNT-FGF activity and then “converted” into posterior NC through the progressive accumulation of neural plate border/definitive NC markers within BMP-responsive cells. The posterior character of these incipient trunk NC precursors and their progeny is likely to be maintained via continuous CDX2 expression and further potentiation of trunk HOX activities as indicated by the expression profiles of these genes in both d6 precursor and d9 trunk NC cells (**Figure 4**, **S4, Tables S2-4**) and their reported roles in trunk NC specification. However, other trunk NC-specific regulators may also be involved in this process and loss-/ gain-of-function approaches are required to dissect their exact involvement in programming trunk identity.

## Acknowledgements

We would like to thank Gabsang Lee and Lorenz Studer for providing the PHOX2B-GFP and SOX10-GFP reporter hPSC lines respectively. T.F. was supported by a University of Sheffield, Biomedical Science Departmental PhD studentship. A.T. is supported by funding from the BBSRC (New Investigator Research Grant, BB/P000444/1), the Royal Society (RG160249) and the Children’s Cancer and Leukaemia Group/Little Princess Trust (CCLGA 2016 01). K.A. and P.W.A. are supported by the EU 7th Framework project PluriMes. M.G. was supported by a BBSRC grant (BB/J015539/1). J.B. is supported by the Francis Crick Institute which receives its funding from Cancer Research UK (FC001051), the UK Medical Research Council (FC001051), and the Wellcome Trust (FC001051). V.W. is supported by an MRC Programme Grant (Mr/K011200/1). We would like to thank Vicki Metzis, Celine Souilhol, Ben Steventon, Matt Towers and Heiko Wurdak for critical reading of the manuscript.

## Author Contributions

Conceptualization: TJRF, AT

Data Curation: IG

Formal Analysis: TJRF, IG, AT

Resources: JB, MG, TJRF, KN, KA, AT

Funding Acquisition: AT, PWA, VW, JB, MG, KA, MRG

Investigation: TJRF, IG, MW, ES, OT, JOSH, DS, PH, MG, AT

Project Administration: AT

Supervision: AT, PWA

Writing – Original Draft Preparation: AT with input from TJRF, IG

Writing – Review & Editing: All co-authors

## Supplemental Figure Legends

**Figure S1.**
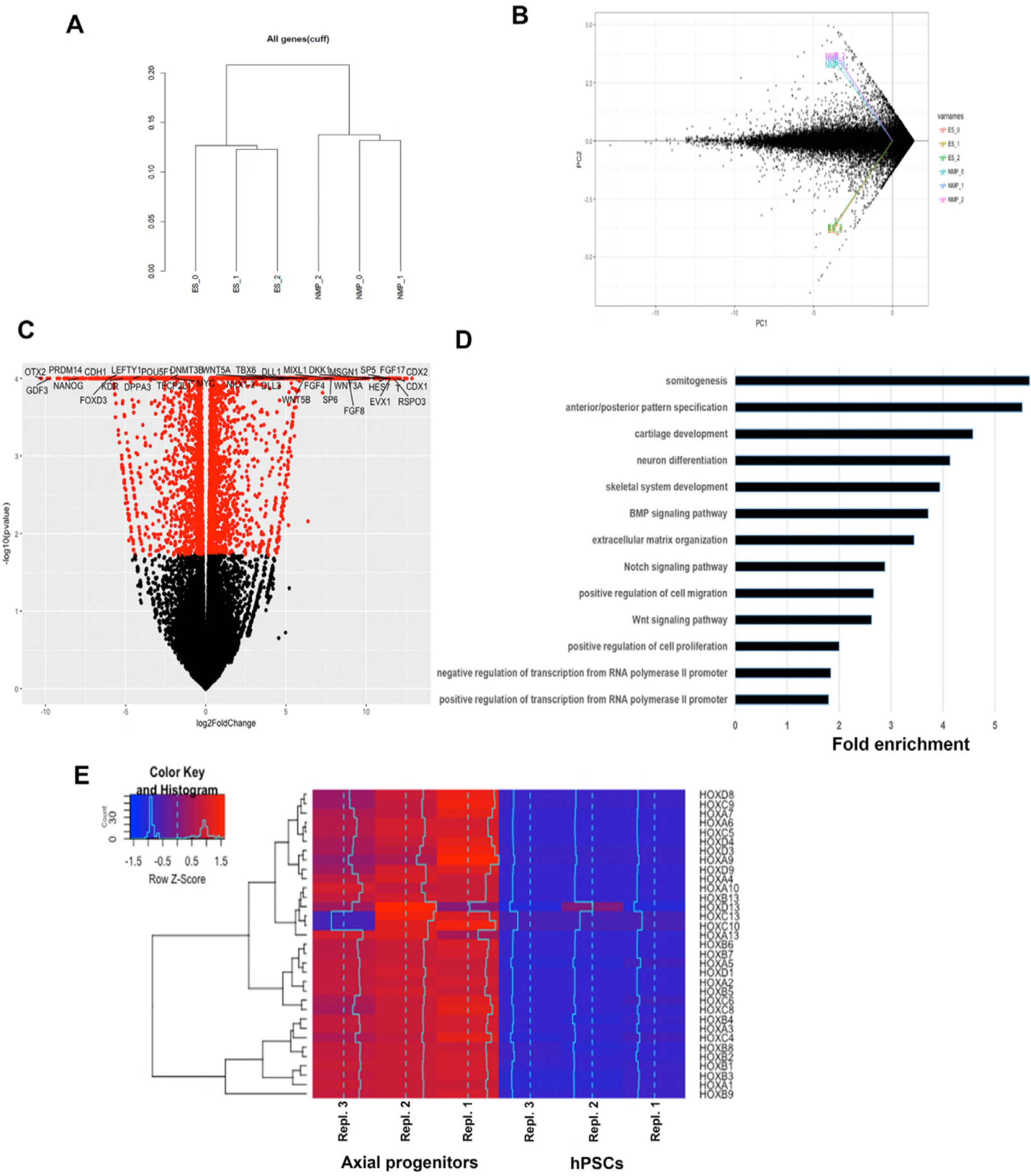
(A) Dendrogram showing the clustering of the individual axial progenitor (NMP) and hESC sample replicates based on RNAseq expression data. (B) Principal Component Analysis (PCA) plot indicates a good separation, in terms of gene expression, between the two conditions and a good replication among samples. ((C) Volcano plot reporting pvalue (y axis) as a function of log2 fold change (x axis) between the axial progenitor and hPSC groups. The red dots depict significant genes (pvalue<=0.05). Top differentially expressed transcriptional regulators of axis elongation and WNT/FGF/NOTCH/RA signalling pathway components are shown. (D) Top gene ontology groups significantly enriched in axial progenitors vs hPSCs. (E) Hierarchical clustering heatmap showing the expression values of HOX genes in axial progenitor and hPSC sample replicates. The expression values (FPKM) were scaled to the row mean. The color key relates the heat map colors to the standard score (z-score).

**Figure S2.**
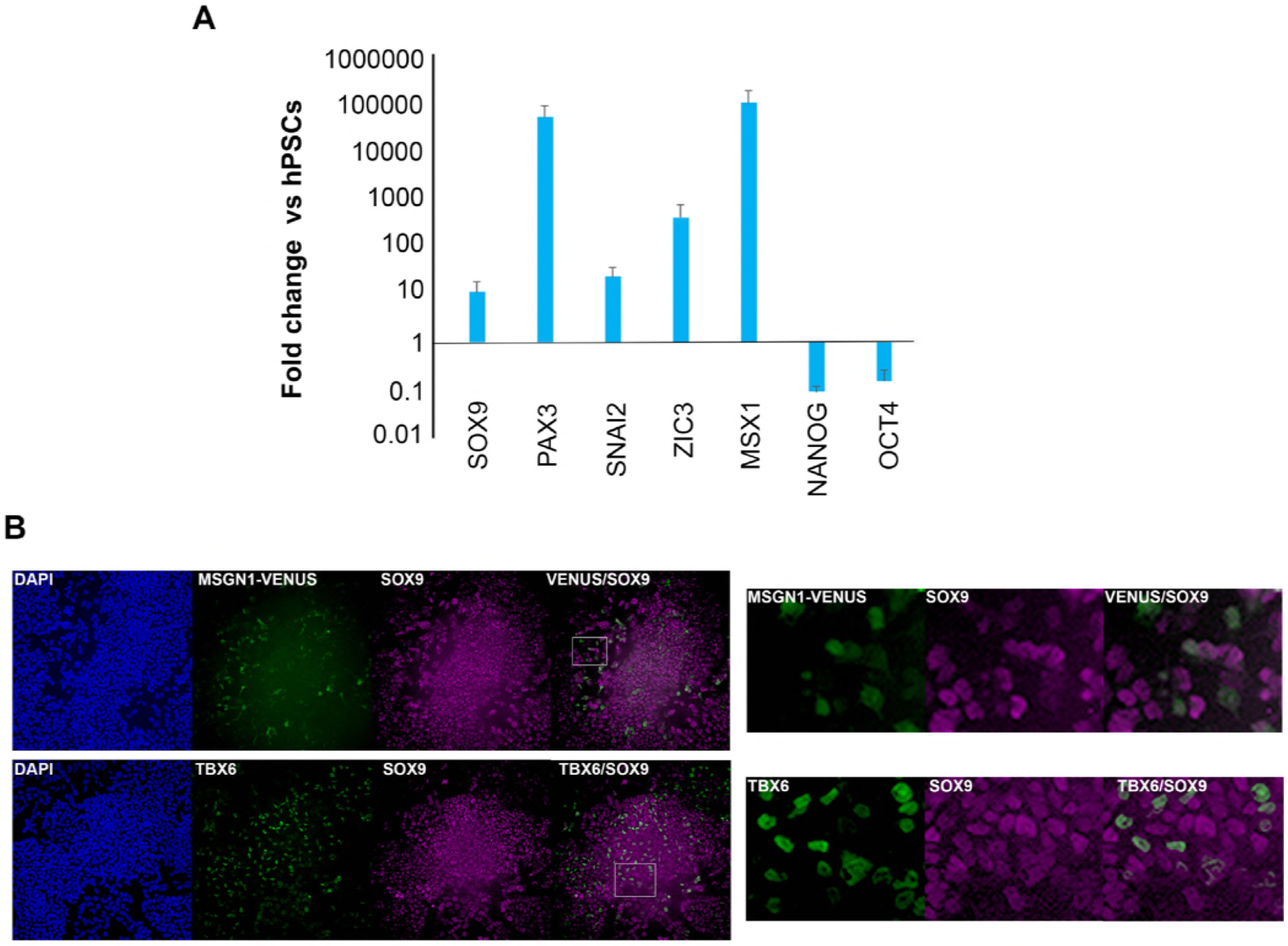
(A) qPCR expression analysis of indicated NC/border and pluripotency markers in axial progenitors vs hPSCs. Error bars=S.E.M. (n=3). (B) Immunofluorescence analysis of SOX9 expression together with MSGN1-VENUS (top) and TBX6 (bottom) in axial progenitors derived from a MSGN1-VENUS or wild type hPSCs respectively.

**Figure S3.**
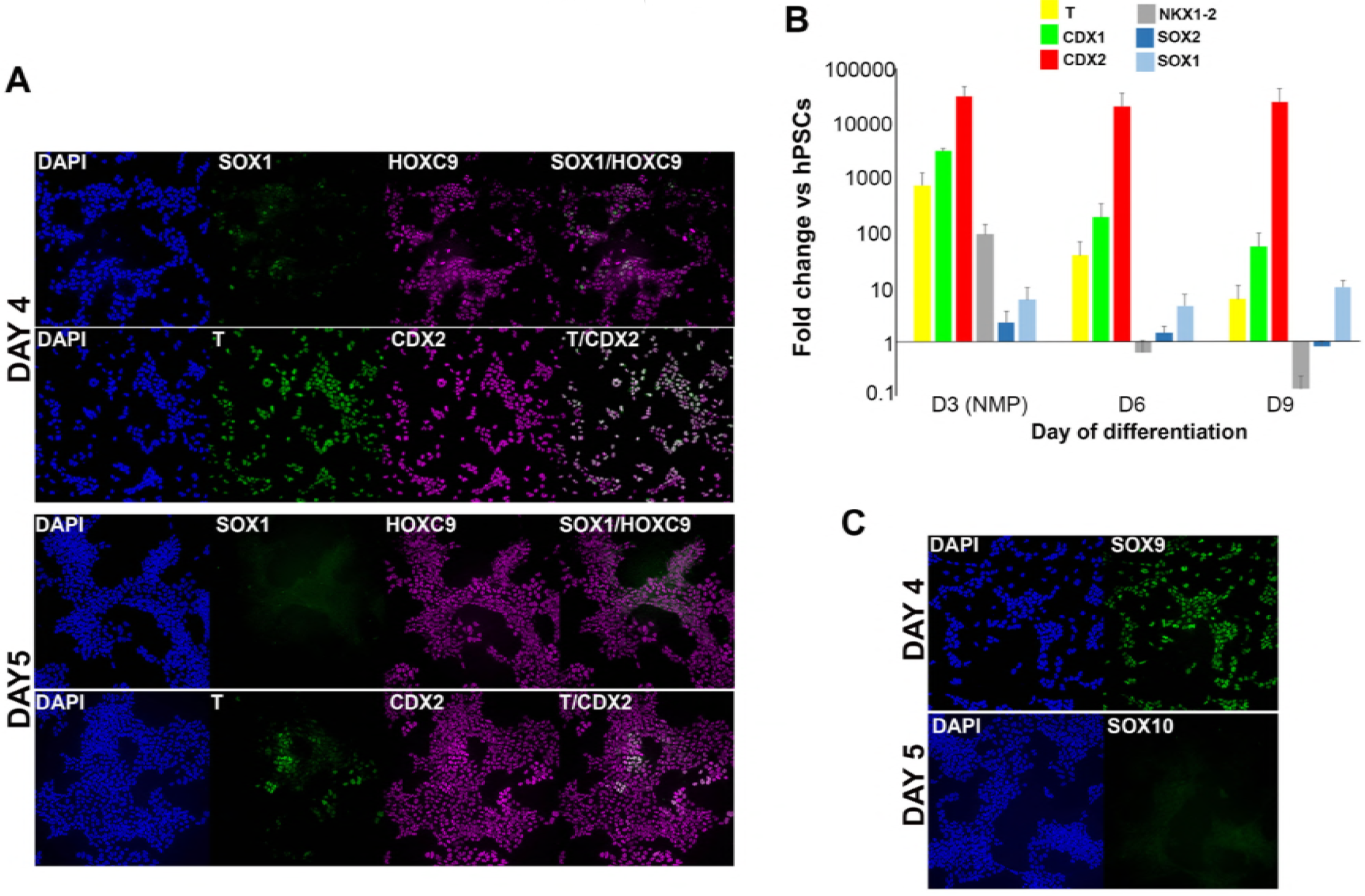
(A) Immunofluorescence analysis of indicated markers one (day 4 of differentiation) or two (day 5 of differentiation) days after re-plating axial progenitors into NC inducing conditions.(B) qPCR expression analysis of indicated axial progenitor (NMP) and neural markers during differentiation toward TNC. Error bars=s.e.m. (n=3) (C) Immunofluorescence analysis of indicated NC markers one (day 4 of differentiation) or two (day 5 of differentiation) days after re-plating axial progenitors into NC inducing conditions.

**Figure S4.**
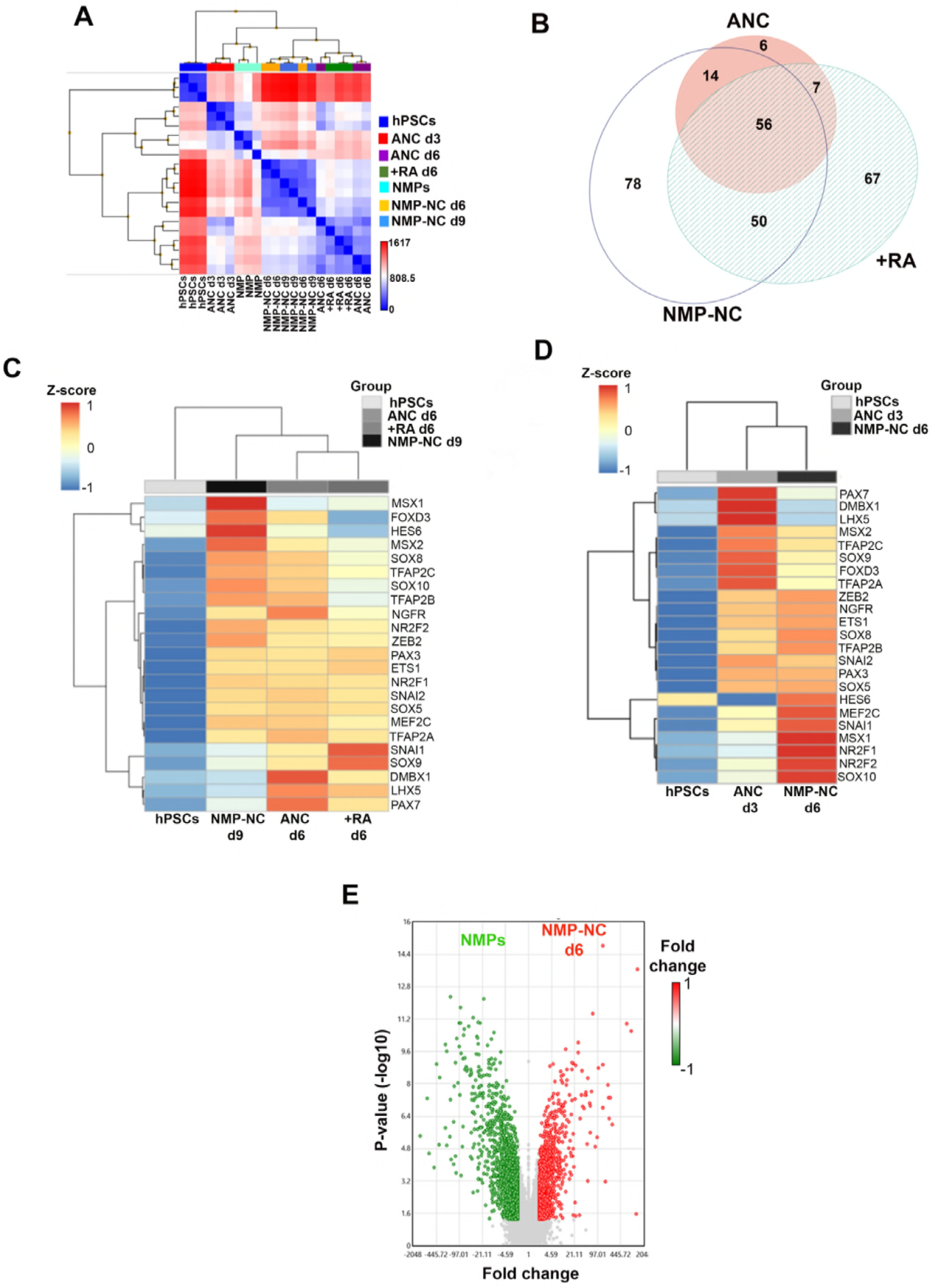
(A) Heatmap showing Pearson’s signal correlation between indicated samples. (B) Venn diagram showing the overlap between transcription factors (TFs) enriched (≥2-fold relative to undifferentiated hPSCs, FDR ≤ 0.05) in each indicated NC group of distinct axial identity. The detailed list of all TFs can be found in Table S3. (C) Heatmap showing relative expression of representative NC markers in NC populations of distinct axial identity and hPSCs. (D) Heatmap showing relative expression of representative NC markers in trunk and anterior cranial NC precursors and hPSCs. (E) Volcano plot depicting upregulated (red) and downregulated (green) genes in d6 trunk NC precursors (NMP-NC) vs axial progenitors (=NMPs).

**Figure S5.**
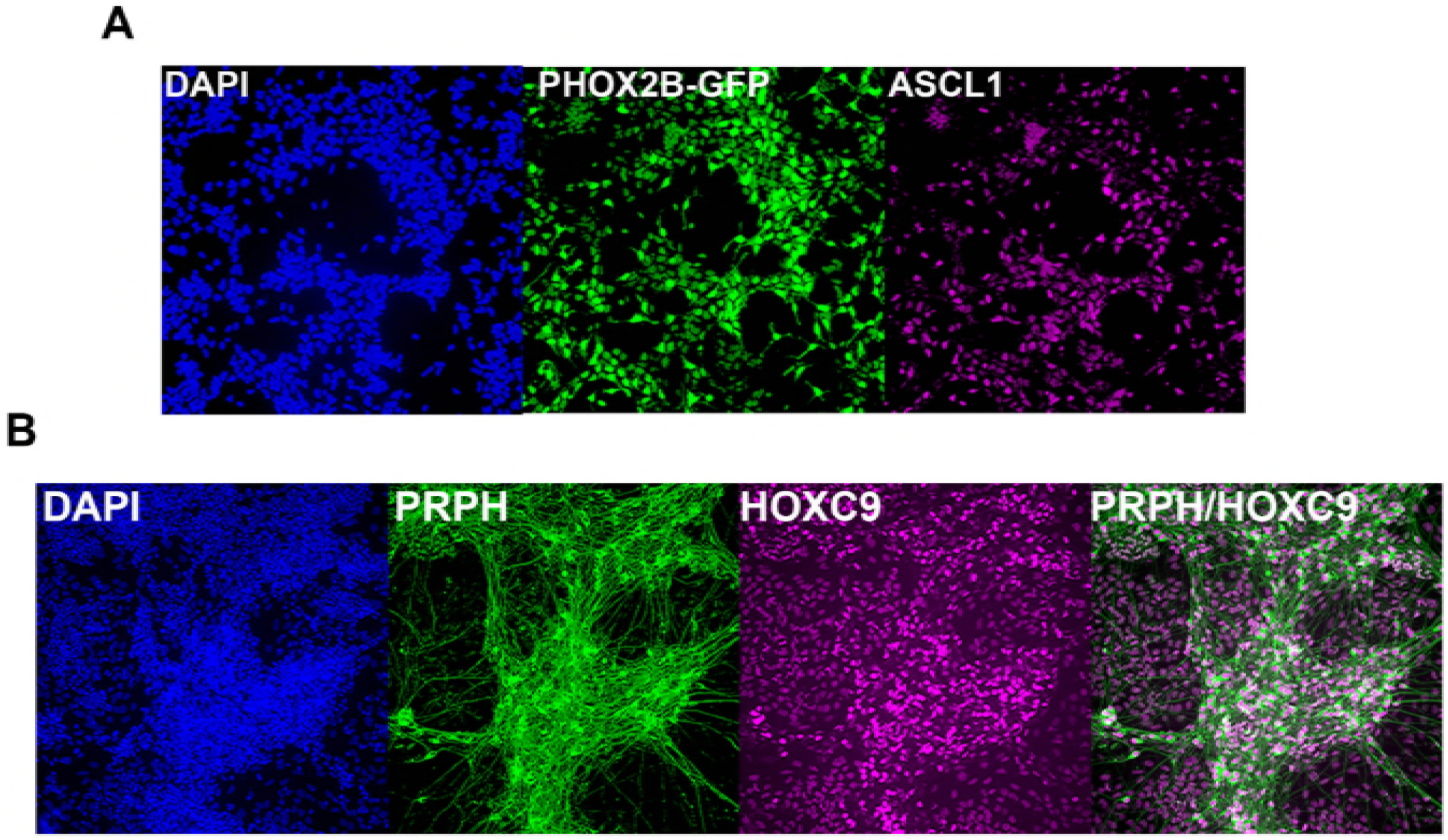
(A) Immunofluorescence analysis of PHOX2B-GFP and ASCL1 expression in d12 SAP cells derived from axial progenitors as shown in Fig. 5A. (B) Immunofluorescence analysis of PRPH and HOXC9 expression in sympathetic neurons derived from axial progenitors as shown in Fig. 5A.

**Table S1.** Significantly up-and downregulated transcripts, GO enrichment and TF signatures from RNAseq analysis

**Table S2.** List of genes upregulated in different NC populations and GO enrichment analysis

**Table S3.** List of transcription factors shared between different NC populations

**Table S4.** List of all genes up-and down-regulated in indicated NC populations and their progenitors.

**Table S5.** List of primers

## METHODS

### Cell culture and differentiation

We employed the following hPSC lines: a Shef4-derived Sox2-GFP reporter hESC line (44), the H9-derived SOX10-GFP (15) and PHOX2B-GFP (18) reporter hESC lines, the MSGN1-VENUS reporter hiPSC line, and the unmodified Mastershef7 hESC line (44) and an iPSC line (MIFF-1) derived from a healthy individual (85). The MSGN1-Venus reporter line was generated by Transposon mediated BAC transgenesis using protocols described by (86). In brief, a human BAC (RP11-12L16) with piggyBac transposon repeats flanking the bacterial backbone and with Venus inserted directly after the initiating methionine of MSGN1 was transfected together with a piggyBac Transposase into NCRM1 iPSCs. Cells were cultured in feeder-free conditions in either Essential 8 (Thermo Fisher) or mTeSR1 (Stem Cell Technologies) medium on laminin 521 (Biolamina) or vitronectin (Thermo Fisher). All differentiation experiments were carried out in at least three different hPSC line. For NMP/axial progenitor differentiation hPSCs were dissociated using PBS/EDTA and plated at a density of 55,000 cells/cm^2^ (density optimised for 12-well plates) on fibronectin (Sigma) or vitronectin (Thermo Fisher)-coated wells, directly into NMP-inducing medium containing CHIR99021 (Tocris), FGF2 (20 ng/ml, Peprotech) and ROCK inhibitor Y-27632 for the first only day (10 μM, Tocris). We observed some variation in terms of induction of T+SOX2+ NMPs both between hPSC lines and also batches of CHIR99021 and thus the concentration of the latter was varied between 3-4 μM. BMP inhibition was carried out using LDN193189 (Tocris) at 100 nM. For trunk NC differentiation day 3 hPSC-derived axial progenitors were dissociated using accutase and re-plated at a density 30,000 cells /cm^2^ on Geltrex (Thermo Fisher)-coated plates directly into NC-inducing medium containing DMEM/F12 (Sigma), 1× N2 supplement (Thermo Fisher), 1× Non-essential amino acids (Thermo Fisher) and 1x Glutamax (Thermo Fisher), the TGFb/Nodal inhibitor SB431542 (2 μM, Tocris), CHIR99021 (1 μM, Tocris), DMH1 (1 μM, Tocris), BMP4 (15 ng/ml, Thermo Fisher) and Y-27632 (10 μM, Tocris). The medium was replaced at days 5 and 7 of differentiation but without the ROCK inhibitor and trunk NC cells were analysed either at day 8 or 9. For cranial neural crest differentiation hPSCs were dissociated using accutase and plated under the same NC-inducing conditions as described above for 5-6 days. For posterior cranial/vagal/cardiac NC generation d4 differentiated anterior NC progenitors induced as described above were treated with retinoic acid (1 μM, Tocris) in the presence of the NC-inducing medium till day 6 of differentiation. For sympathoadrenal progenitor (SAP) differentiation d8 trunk NC cells were re-suspended at a density of 200-300,000 cells /cm^2^ on Geltrex (Thermo Fisher)-coated plates directly into medium containing BrainPhys neuronal medium (Stem Cell Technologies), 1× B27 supplement (Thermo Fisher), 1x N2 supplement (Thermo Fisher), 1× Non-essential amino acids (Thermo Fisher) and 1× Glutamax (Thermo Fisher), BMP4 (50 ng/ml, Thermo Fisher), recombinant SHH (C25II) (50 ng/ml, R&D) and purmorphamine (1.25-1.5 μM, Millipore) and cultured for 4 days (d12 of differentiation). For further sympathetic neuron differentiation d12 SAP cells were switched into a medium containing BrainPhys neuronal medium (Stem Cell Technologies), 1× B27 supplement (Thermo Fisher), 1× N2 supplement (Thermo Fisher), 1× Non-essential amino acids (Thermo Fisher) and 1x Glutamax (Thermo Fisher), ascorbic acid (200 μM, Sigma), NGF (10 ng/ml, Peprotech), BDNF (10 ng/ml, Peprotech) and GDNF (10 ng/ml, Peprotech) and cultured for a further 6-8 days. For paraxial mesoderm differentiation d3 axial progenitor cultures were treated with accutase and replated at a density of 45,000/cm^2^ on 12-well Geltrex-coated plates in N2B27 containing FGF2 (40 ng/ml, Peprotech) and CHIR99021 (8 μM, Tocris) for two days. For neural differentiation d3 axial progenitor cultures were treated with accutase and replated at a density of 45,000/cm^2^ on 12-well Geltrex-coated plates in N2B27 containing 100 nM retinoic acid (Tocris) for 2-3 days.

### RNA sequencing

#### Sample preparation

For RNA sequencing we employed hESCs or axial progenitors following culture on fibronectin in FGF2 (20 ng/ml) and CHIR99021 (3 μM). Total RNA from NMPs and hESCs was harvested using the RNeasy kit (Qiagen) according to the manufacturer’s instructions.

#### Library preparation/sequencing

Total RNA was processed according to the TruSeq protocol (Illumina). Three separate RNA libraries (biological replicates) were barcoded and prepared for hPSCs and D3 axial progenitors. Library size, purity and concentration were determined using the Agilent Technologies 2100 Bioanalyzer. For sequencing, four samples were loaded per lane on an Illumina Genome Analyzer Hiseq2500.

#### RNAseq quality control and mapping

The quality of raw reads in fastq format was analyzed by FastQC (http://www.bioinformatics.babraham.ac.uk/projects/fastqc). Adapter contamination and poor quality ends were removed using Trim Galore v. 0.4.0 (Babraham Bioinformatics-Trim Galore! Available at: http://www.bioinformatics.babraham.ac.uk/projects/trim_galore/). Single-end clean reads were aligned to the human reference genome (hg38 assembly) using Tophat2 v2.0.13 (87).

#### RNA seq data analysis

Read alignments were sorted with SAMtools v1.1 before being counted to genomic features by HTSeq version 0.6.0 (88). The average overall read alignment rate across all samples was 94.3%. Differential gene expression was performed using DESeq2 version 1.16.1 (89). in R version 3.3.3. Genes were considered significantly differentially expressed (DE) with a Benjamini-Hochberg adjusted pvalue <=0.05 and a log2FoldChange> |1|. Gene Ontology (GO) biological processes (BP) enrichment analysis was carried out for DE genes using the DAVID gene ontology functional annotation tool (https://david.ncifcrf.gov/) (90, 91) with default parameters. We considered as significant terms having a FDR adjusted pvalue <=0.05, which is derived from a modified Fisher’s exact test.

### Microarrays

#### Sample preparation and processing

Samples were prepared according to the Affymetrix WT Plus protocol for Gene Chip ^®^ Whole Transcript Expression Arrays. Briefly 200ng of high quality total RNA, (RNA integrity number (RIN) greater than 9), was converted to double stranded cDNA with the introduction of a T7 polymerase binding site. This allowed the synthesis of an antisense RNA molecule against which a sense DNA strand was prepared. The RNA strand was digested and the resulting single stranded DNA fragmented and biotin labelled. Along with appropriate controls the labelled fragmented DNA was hybridised to Affymetrix Clariom D arrays overnight using the Affymetrix 640 hybridisation oven; 16 hours with rotation at 60 rpm at 45°C. The arrays were washed and stained according to standard protocols which allowed the introduction of streptavidin-phycoerythrin in order to generate a fluorescent signal from the hybridised biotinylated fragments. The washed and stained arrays were scanned using the Affymetrix 3000 7G scanner with autoloader. The generated CEL files were taken forward for analysis.

#### Data analysis

Data were analysed using the Affymetrix Transcriptome Analysis Console 4.0 software. Analysis of Expression (Gene+Exon) was used to generate lists of all differentially expressed genes showing >2; <-2 fold Log Change and P<0.05. For the distance matrix (**Figure S4A**), Exploratory Grouping analysis was used. Log_2_ normalised intensity data values were mapped in R using the package ‘pheatmap’ with correlation clustering by gene. Gene ontology analysis was carried out using the ToppGene suite (https://toppgene.cchmc.org/enrichment.jsp) (92). Area proportional 3-Venn diagrams were drawn using the eulerApe software (93).

### Quantitative real time PCR

Total RNA from different samples was harvested using the RNeasy kit (Qiagen) according to the manufacturer’s instructions and digested with DNase I (Qiagen) to remove genomic DNA. First strand cDNA synthesis was performed using the Superscript III system (Thermo Fisher) using random primers. Quantitative real time PCR was carried out using the Applied Biosystems QuantStudio™ 12K Flex thermocycler together with the Roche UPL system. Primer sequences are shown in Table S5.

### Flow cytometry

Cells were lifted into a single cell suspension using Accutase (as previously described) and resuspended in FACS buffer (DMEM with 10% v/v FCS) to neutralise Accutase before centrifugation at 1100rpm/4min. Cells were then resuspended in FACs buffer at 1×106 cells/ml and 100,000 cells were placed in FACs tubes (Falcon 352053). A GFP baseline was set using unmodified wild type control cells.

### Immunofluorescence

Cells were fixed for 10 minutes at 4°C in 4% paraformaldehyde (PFA) in phosphate buffer saline (PBS), then washed in PBST (PBS with 0.1% Triton X-100) and treated with 0.5 M glycine/PBST to quench the PFA. Blocking was then carried for 1-3 h in PBST supplemented with 3% donkey serum/1% BSA at room temperature or overnight at 4μC. Primary and secondary antibodies were diluted in PBST containing in PBST supplemented with 3% donkey serum/1% BSA. Cells were incubated with primary antibodies overnight at 4°C and with secondary antibodies at room temperature for 2 h in the dark. Cell nuclei were counterstained using Hoechst 33342 (1:1000, Thermo Fisher) and fluorescent images were taken using the InCell Analyser 2500 system (GE Healthcare). We used the following antibodies: anti-T (1:200; AF2085, R&D), anti-SOX2 (1:200; ab92494, Abcam), anti-SOX9 (1:200; 82630, CST), anti-SNAI2 (1:400; C19G7, CST), anti-PAX3 (1:50; DSHB), anti-phosphoSMAD1/5/9 (1:100; D5B10, CST), anti-SOX10 (1:200; D5V9L, CST), anti-SOX1 (1:100; AF3369, R&D), anti-TBX6 (1:50, AF4744, R7D) anti-TH (1:1000; T1299, SIGMA), HOXC9 (1:50; ab50839, Abcam) anti-PRPH (1:100; AB1530, Millipore), anti-CDX2 (1:200; ab76541, Abcam), anti-ASCL1 (1:100; 556604, BD Pharmigen), anti-GATA3 (1:100; sc-269, Santa Cruz), anti-GFP (1:1000; ab13970, Abcam), anti-ISL1 (1:100, DSHB), anti-PHOX2B (1:100; sc-376997, Santa Cruz). Images were processed using Photoshop and Fiji. Nuclear segmentation followed by single cell fluorescence quantification was performed as described previously using Fiji and the MultiCell3D application as described previously (35, 94).

#### Data availability

The microarray and RNAseq data have been deposited to GEO (GSE109267 and GSE110608).

